# Chromosomally-Encoded Hok/Sok Toxin/Antitoxin Systems Inhibit Phage Infection

**DOI:** 10.64898/2026.11.02.763616

**Authors:** Eunsol Seo, Maura E. Cingle, Thomas K. Wood

## Abstract

The primary physiological role of toxin/antitoxin (TA) systems is phage defense; they are also used to stabilize mobile genetic elements. In the initial publication in 1996, plasmid-encoded Hok/Sok, a type I TA system (protein toxin, RNA antitoxin that inhibits toxin mRNA), was shown to inhibit T4 phage infection in *Escherichia coli* by reducing its efficiency of plating and burst size and by increasing the time for mature phage to form (latent period). This publication also identified that the mechanism of Hok/Sok inhibiting T4 was host transcription shutoff, arguably the most-general phage inhibition system since most phages impair host transcription. However, recently, the role of Hok/Sok in phage inhibition has been challenged. Therefore, here we investigated the role of four *E. coli* chromosomally-expressed, *hok*^+^*/sok*^+^ loci and the one orphan *hok*^+^ locus and found that the Hok proteins inhibit the infection of *E. coli* by BASEL phage 48, a Vequintavirus with myovirus morphology. Of the five Hok proteins, the greatest phage inhibition was by HokB, and infection by BASEL phages 34 and 39 were also inhibited by HokB. We also find BASEL phage 48 requires the WecB-generated receptor NGR for infection. Therefore, chromosomally-derived orphan Hok proteins, produced at physiological levels, are relevant for phage inhibition.

## INTRODUCTION

Almost all bacteria contain toxin/antitoxin (TA) systems, and these loci are most-commonly linked to mobile element stabilization and phage defense (Song and Wood, 2020). Based on how the antitoxin masks toxin activity as RNA or as a binding protein/enzyme, TA system are classified into nine types (Seo et al., 2026a). In general, toxins function by inhibiting DNA replication and cell division, or by degrading mRNAs/tRNAs and energetic molecules such at ATP (Jurėnas et al., 2022).

The type I system Hok/Sok was the first TA system shown to inhibit phage infection (Pecota and Wood, 1996). Specifically, *hok*^*+*^*/sok*^*+*^ was found to inhibit consistently T4 phage in three ways at three different multiplicity of infections (MOI’s, 0.001, 0.01, and 0.12): (i) the absence of *hok/sok* increased sensitivity to T4, (ii) overexpression of Hok/Sok rendered the cell insensitive to T4, and (iii) the wild-type plasmid R1, which contains *hok*^*+*^*/sok*^*+*^ with natural expression, reduced sensitivity to T4 phage (Pecota and Wood, 1996). Furthermore, it was determined that Hok/Sok inhibited T4 phage infection by reducing its efficiency of plating and burst size and by increasing the time for mature phage to form (latent period). Since T4 phage stops transcription in 3 to 4 min and Sok antitoxin is labile (∼ 1 min), the mechanism for *hok*^*+*^*/sok*^*+*^ inhibition of T4 phage was found to be transcription shutoff by T4 (Pecota and Wood, 1996). In addition, lambda-virulent and T5 phage were slightly inhibited by *hok*^*+*^*/sok*^*+*^ at three MOI’s (Pecota and Wood, 1996), whereas there was no inhibition with T1 and T7 phage for *hok*^*+*^*/sok*^*+*^ (Pecota and Wood, 1996). Hence, *hok*^*+*^*/sok*^*+*^ from plasmid R1 was found to provide robust phage inhibition, and the paradigm of TA systems inhibiting phages was established (Song and Wood, 2020).

Cementing the paradigm of phage inhibition by TA systems, next the type II MazE/MazF (Hazan and Engelberg-Kulka, 2004), and the type III ToxN/ToxI (Fineran et al., 2009) TA systems were found to inhibit phage infection. Recently, the paradigm has been extended to nearly all TA systems with the type IV ShosT/ShosA (Pu et al., 2025), the type VII systems PfkAB/PfpC (Guo et al., 2024) and Hha/TomB (Seo et al., 2026b), and the type IX CloA/CloB (Clover) (Yamaguchi et al., 2025). However, there is no report chromosomal copies of *hok*^*+*^*/sok*^*+*^ and phage inhibition.

In *E. coli* K-12 MG1655, there are five *hok*^*+*^ genes; all of which were deemed inactive (Pedersen and Gerdes, 1999). Of these, (i) *hokA*^*+*^, *hokB*^*+*^, *hokC*^*+*^, and *hokE*^*+*^ have all of the regulatory elements of the canonical *hok*^*+*^*/sok*^*+*^ of R1, (ii) *hokA*^*+*^*/sokA*^*+*^, *hokC*^*+*^*/sokcC*^*+*^, and *hokE*^*+*^*/sokE*^*+*^ contain insertion sequences (but not within *hok*^*+*^), and (iii) *hokD*^*+*^ is transcribed as part of the *relBE*^*+*^ TA locus (Pedersen and Gerdes, 1999). Moreover, toxins HokA, HokB, HokC, and HokD are active as toxins (Pedersen and Gerdes, 1999).

Subsequently, HokB was linked to persistence through GTPase Obg (Verstraeten et al., 2015), but there was no persister phenotype for the *hokB* mutation and the results were based on non-physiological levels of HokB from overexpression from plasmids (Wood and Song, 2020). Furthermore, HokB mRNA has been shown to be edited by adenosine deaminase TadA, resulting in a tyrosine to cysteine substitution at amino acid 29 that forms an additional disulfide bond in the periplasm and increases toxicity (Didi et al., 2025).

Here, we investigate whether *hok*^*+*^*/sok*^*+*^ of R1plasmid-based paradigm of phage inhibition also applies to chromosomal copies of the *E* .*coli* K-12 *hok*^*+*^*/sok*^*+*^ loci. We find four of the five *hok*^*+*^ chromosomal loci serve to inhibit the infection of BASEL phage 48 (*Vequintavirus* 137.6 kb) and that HokB also inhibits infection by BASEL phages 34 and 39.

## RESULTS

### Rationale

Given that plasmid-based *hok*^*+*^*/sok*^*+*^ inhibits phage T4 phage (Pecota and Wood, 1996), we investigated whether the five chromosomal copies of *hok*^*+*^ also inhibit phages. We focused initially on phage 48 of the BASEL collection of 69 *Escherichia coli* lytic phages (Maffei et al., 2021) since we recently found it is affected by the type VII TA system Hha/TomB (Seo et al., 2026b).

### Chromosomal *hokABCDE* screen with BASEL phage 48

Using the isogenic, single-gene mutants of the Keio Collection (Baba et al., 2006), we found that deleting each of the five chromosomally-encoded *hok* loci had a substantial effect on phage infection (**Fig. 1ABC**). We found consistently at three different BASEL phage 48 titres (MOI 0.1, 0.01, and 0.001) that four of the five *hok* deletions reduced the ability the cell to combat BASEL phage 48. Moreover, we found the *hokB* deletion has the largest effect at each of the three MOI’s (**Fig. 1ABC**). As a negative control, we found that all five of the *hokABCDE* deletions grew at almost identical rates in the absence of phage (**Fig. 1D**). Hence, *hokABCDE* are active in phage defense and their impact is not related to growth defects.

**Fig. 1.**
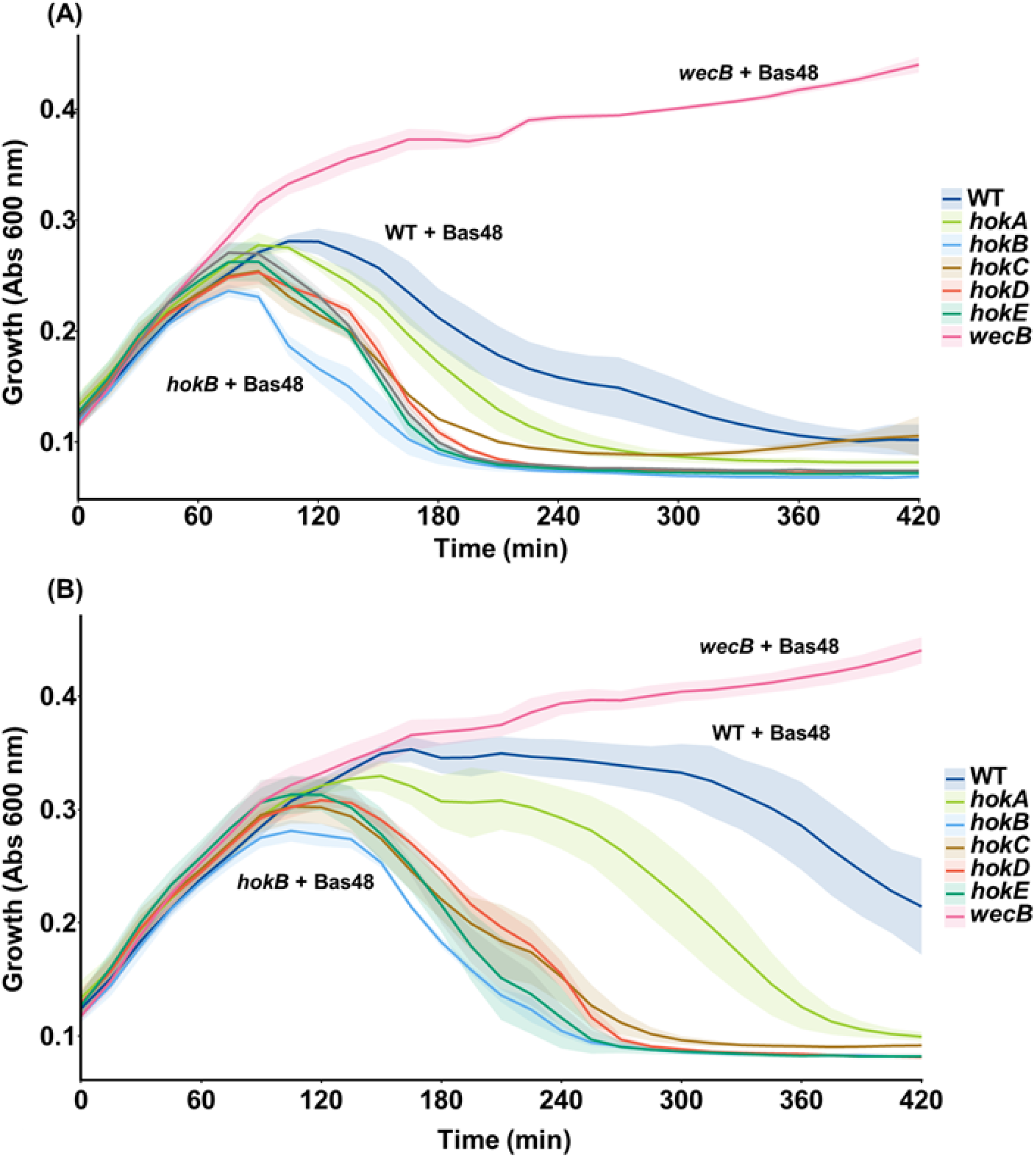
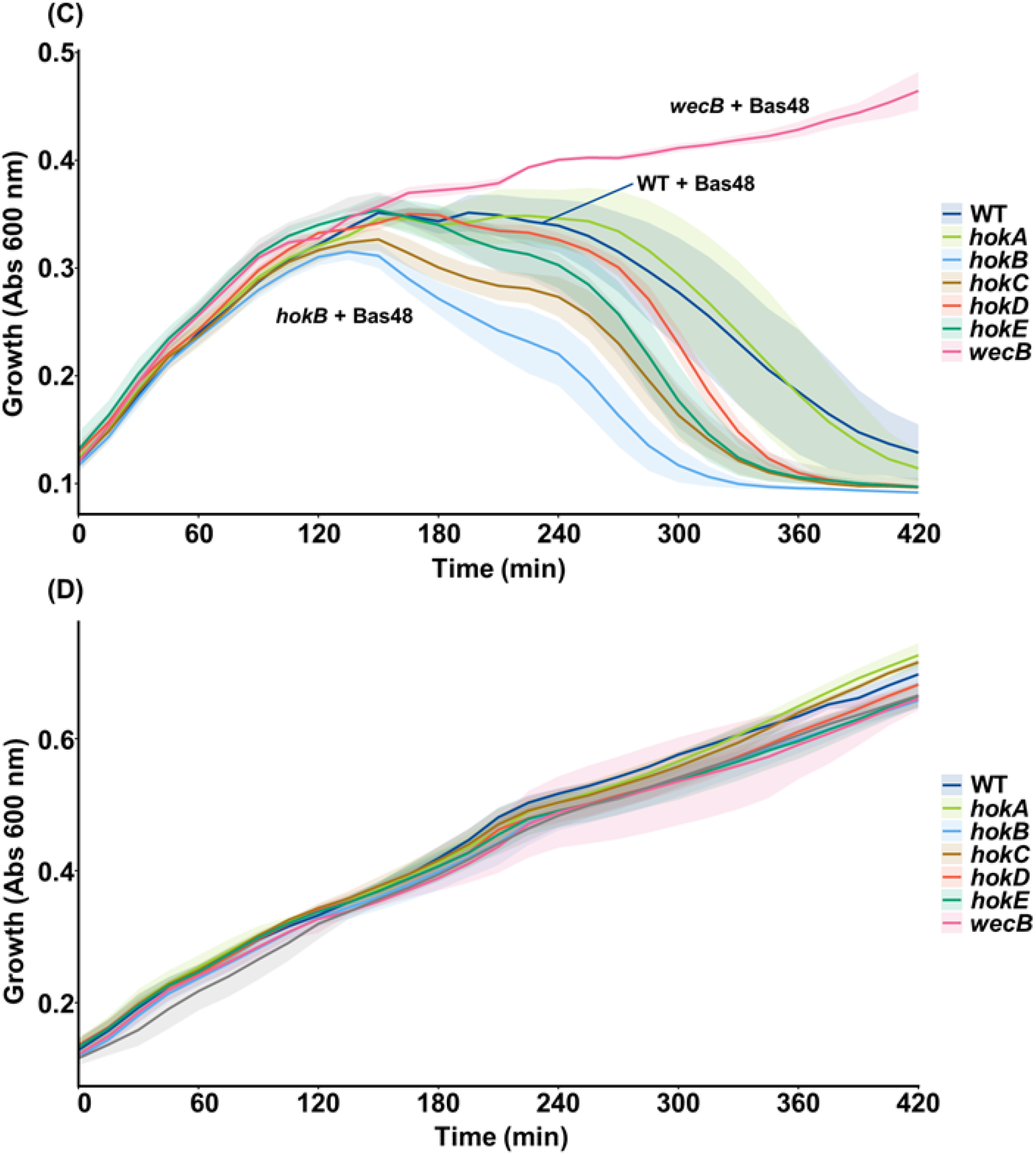
Chromosomally-encoded Hok proteins inhibit BASEL phage 48 infection. *hokA, hokB, hokC, hokD*, and *hokE* deletions with BASEL phage 48 at 0.1 MOI (**A**), *hokABCDE* deletions with BASEL phage 48 at 0.01 MOI (**B**), and *hokABCDE* deletions with BASEL phage 48 at 0.001 MOI (**C**). BASEL phage 48 was added to cells grown to a turbidity of 0.1 at 600 nm. Growth was recorded in a microplate Tecan reader with some shaking. **(D)** The *hokABCDE* deletions do not affect growth in the absence of phage. Results from three independent cultures with standard deviations shown as shading.

### WecB is the likely surface receptor for BASEL phage 48

We also hypothesized that WecB, a cytoplasmic UDP-*N-*acetylglucosamine 2-epimerase, would be required as the receptor for BASEL phage 48. This is based on the report that the *N*-acetylmannosamine-based carbohydrate polymer novel surface glycan receptor is exported to the cell surface of *E. coli* and is the likely receptor for *E. coli* phage N4, a member of the *Enquatrovirus g*enus, as well as is the likely receptor for the *Vequintavirinae* subfamily phages (Sellner et al., 2021), that includes BASEL phage 48. In agreement with our hypothesis, we found the *wecB* isogenic mutation rendered the cell completely resistant to BASEL phage 48 at all three MOI’s (**Fig. 1ABC**). Moreover, the *wecB* mutation had no effect on growth in the absence of phage (**Fig. 1D**).

Hence, WecB is required for BASEL phage 48 infection, and the *wecB* mutation serves well as a negative control for the evaluation of the effect of Hok and phage infection.

### HokB is also inhibits BASEL phages 34 and 39

Since the *hokB* mutation had the greatest impact of BASEL phage 48 infection (**Fig. 1ABC**), we focused on HokB. To investigate if HokB could inhibit other 80 BASEL phages, we tested BASEL phages 34 (siphovirus) and 39 (myovirus) with the Δ*hokB* strain. We found that upon deleting *hokB*, the cell became much more sensitive to BASEL phage 39 and somewhat more sensitive to BASEL phage 34 (**Fig. 2**).

**Fig. 2.**
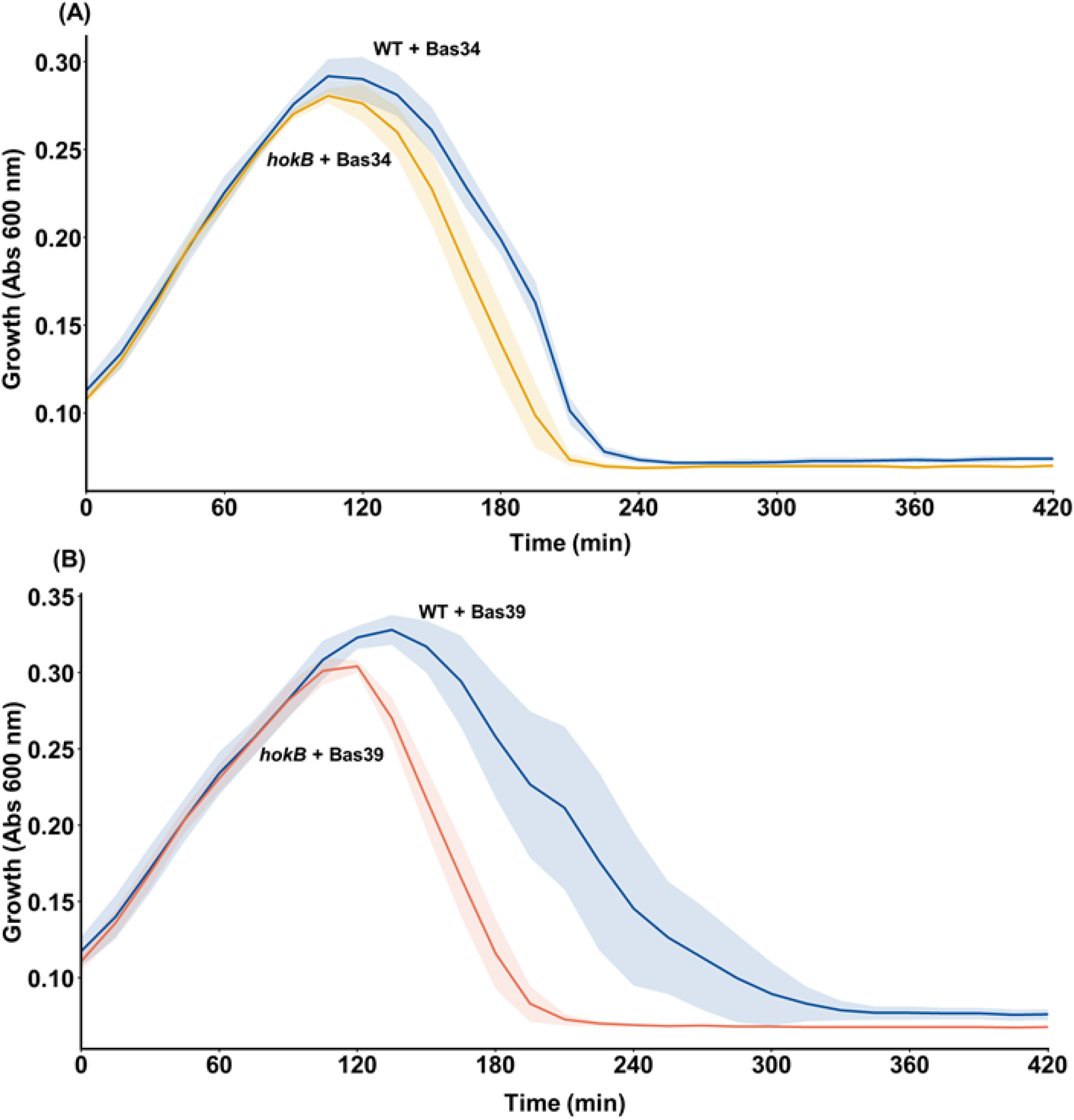
Chromosomally-encoded HokB inhibits BASEL phages 34 and 39. *hokB* deletion with BASEL phage 34 (**A**), and *hokB* deletion with BASEL phage 39 (**B**). BASEL phages at 0.1 MOI were added to cells grown to a turbidity of 0.1 at 600 nm. Growth was recorded in a microplate Tecan reader with some shaking. Results from three independent cultures with standard deviations shown as shading.

### HokB is active

Using the *hokB* host so that that there is no chromosomal HokB production, we confirmed that HokB is active as a toxin as induction of *hokB* results in both the cessation of growth and loss of viable 85 cells (**Fig. 3AB**). As expected, greater toxicity was seen with increased induction of the plasmid promoter with 1 mM IPTG vs. 0.1 mM IPTG (**Fig. 3B**).

**Fig. 3.**
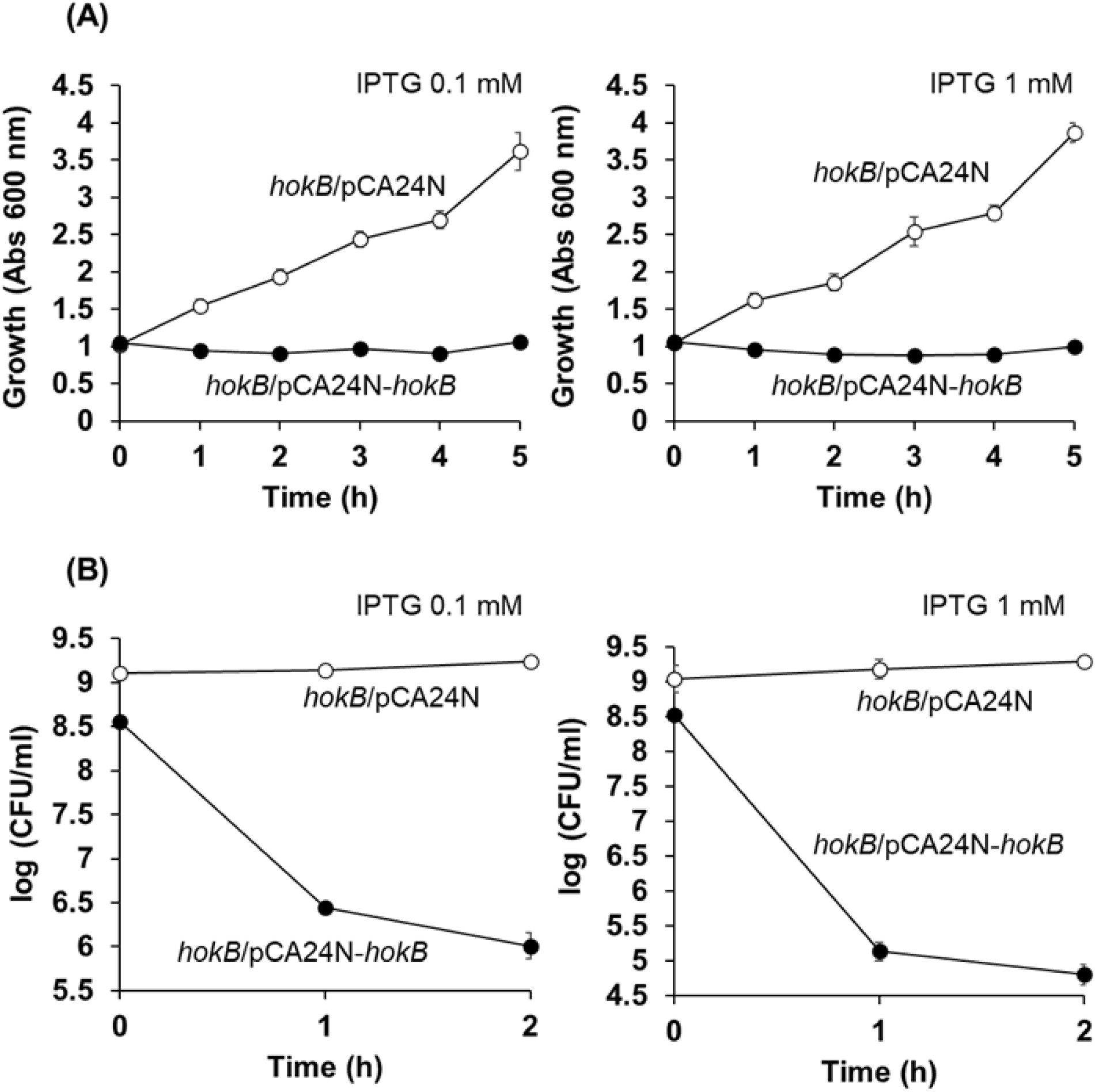
HokB is active. Addition of both 0.1 and 1.0 mM IPTG at time 0 to induce *hokB* from pCA24N-*hokB* in BW25113 results in cessation of growth (**A**) and cell death **(B**). Results from three independent cultures with standard deviations shown as bars.

## DISCUSSION

In this report, we show clearly that the chromosomally-encoded toxin Hha plays a role in phage efense for at least three phages, BASEL phages 48, 34, and 39. We also found that the WecB-generated 90 receptor NGR is the likely infection receptor for BASEL phage 48. Our results provide additional evidence of the relationship for Hok/Sok and phage defense to that of the plasmid-based system used for the initial discovery linking TA system and phage defense (Pecota and Wood, 1996).

Critically, our results are in stark contrast to those of Messineo et al. (Messineo et al., 2026), whoclaimed *hok*^*+*^*/sok*^*+*^ is not involved in phage defense. We surmise that the difference between our results and theirs is that Messineo et al. failed to use an active Hok protein (no Hok activity was reported), since TA system toxins may be inactivated within hours when induced (Fernandez-Garcia et al., 2019).

The most-likely mechanism for Hok/Sok inhibition of phages remains transcription shutoff (Pecota and Wood, 1996). After a phage infects *E. coli*, host transcription is indubitably halted, which leads to cessation in the production of Hok and Sok RNA. Since Sok RNA is more labile than Hok RNA (Gerdes 100 et al., 1986), Hok RNA is translated, leading to membrane disruption. This membrane disruption leads to a reduction in protein production and may lead to the cell entering a persister state (Fernández-García et al., 2024a), which allows phage defense systems like type IV restriction systems to eliminate phage (Fernández-García et al., 2024b).

## METHODS

### Bacterial strains, phages, plasmids, and growth conditions

The bacterial strains, isogenic mutants, and plasmids are described in **Table 1**; we used only *E. coli* K-12 BW25113 and its isogenic mutants. Lytic phages from the BASEL collection were obtained from Alexander Harms (Maffei et al., 2021). Bacteria were grown in lysogeny broth (LB) containing 10 g/L tryptone, 5 g/L yeast extract, and 10 g/L NaCl at 37°C with shaking at 250 rpm. Where applicable, knockout strains were precultured with 50 µg/mL 110 kanamycin. Strains carrying pCA24N-derived plasmids were cultured with 30 µg/mL chloramphenicol forplasmid maintenance.

**Table 1.** *Escherichia coli* strains and plasmids used in this study. Km^R^, kanamycin resistance. Cm^R^, chloramphenicol resistance.

| **Name** | **Genotype** | **Source** |
|---|---|---|
| BW25113 | K-12 *rrnB3* Δ*lacZ4787 hsdR514* Δ*(araBAD*)567 Δ*(rhaBAD*)568 *rph-1* | (Baba et al., 2006) |
| BW25113 *hokA* | Δ*hokA* Ω Km<sup>R</sup> | (Baba et al., 2006) |
| BW25113 *hokB* | Δ*hokB* Ω Km<sup>R</sup> | (Baba et al., 2006) |
| BW25113 *hokC* | Δ*hokC* Ω Km<sup>R</sup> | (Baba et al., 2006) |
| BW25113 *hokD* | Δ*hokD* Ω Km<sup>R</sup> | (Baba et al., 2006) |
| BW25113 *hokE* | Δ*hokE* Ω Km<sup>R</sup> | (Baba et al., 2006) |
| **Plasmidse** |  |  |
| pCA24N | Cm<sup>R</sup>; *lacI*<sup>q</sup> | (Kitagawa et al., 2005) |
| pCA24N-*hokB* | Cm<sup>R</sup>; *lacI*<sup>q</sup> P<sub>T5-lac</sub>::*hokB*<sup>+</sup> | (Kitagawa et al., 2005) |

### Temporal phage assay

Overnight cultures prepared in LB supplemented with the appropriate antibiotics, where required, were used to inoculate antibiotic-free LB at a 1:100 dilution. Once the cultures reached an optical density at 600 nm of approximately 0.1, they were dispensed into 96-well plates. BASEL phages were then added at a multiplicity of infection (MOI) of 0.1. Bacterial growth was monitored using a Tecan microplate reader with intermittent shaking.

## ACKNOWLEDGEMENTS

This work was supported by funds derived from NSF grant 2515448 (UEI: NPM2J7MSCF61). We thank Alexander Harms for providing the BASEL collection of *Escherichia coli* phages.

